# Cell-type Specific Learning of Attentional Gating in Primate Striatum

**DOI:** 10.1101/837740

**Authors:** Kianoush Banaie Boroujeni, Mariann Oemisch, Seyed Alireza Hassani, Thilo Womelsdorf

## Abstract

Cognitive flexibility depends on a fast neural learning mechanism for enhancing momentary relevant over irrelevant information. A possible neural mechanism realizing this enhancement uses fast-spiking interneurons (FSIs) in the striatum to train striatal projection neurons to gate relevant and suppress distracting cortical inputs. We found support for such a mechanism in nonhuman primates during the flexible adjustment of visual attention. FSIs gated visual attention cues during feature-based learning. One FSI population showed stronger inhibition during learning, while another FSI subpopulation showed weaker inhibition after learning signifying post-learning disinhibition. Additionally, a smaller neural subpopulation increased activity when salient distractor events were successfully suppressed. These findings highlight that fast behavioral learning of feature relevance is accompanied by fast neural learning of cell-type specific cortico-striatal gating.

---

Adaptive behavior depends on neural mechanisms that learn which objects in our environment have value and are relevant to achieve our goals. Knowing the value of objects is essential to not be distracted and allocate attention efficiently. The fast learning of object values is enabled by neural circuits in the head of the caudate nucleus in the anterior striatum of the nonhuman primate brain [1–8]. Caudate neurons begin to fire stronger when objects appear at recently rewarded locations [2, 8, 9], and when attention is directed to the rewarded and away from nonrewarded stimuli [7, 10]. Electrical microstimulation suggests that these neuronal activity changes in the caudate are part of a causal processing chain for enhancing reward sensitivity and visuo-motor associations [11–13], as well as for biasing choice and attention [9]. Electrically stimulating the caudate nucleus during the presentation of a colored visual stimulus can enhance the preference to choose that stimulus after stimulation [9]. This color-based choice bias can be independent of the stimulus location, suggesting that caudate learning can change the synaptic gating of cortical inputs in a feature-specific way [7, 10].

Fast learning in the anterior striatum might involve dopaminergic modulation of synaptic weights among neurons receiving updates about the value of visual objects during learning [7, 14]. In the striatum these neurons are spiny projection neurons (SPNs) and recent in-vivo evidence in rodent striatum supports the view that synaptic connections among SPN’s are trained by inhibition from fast spiking interneurons (FSI’s) [15–17]. According to this model cognitive flexibility is achieved through selective FSI-mediated inhibition of SPN neurons processing cortical inputs about distractors and disinhibition of SPNs processing reward predictive, relevant information [18].

We set out to test the key predictions of this model in the nonhuman primate anterior striatum. Our color-based reversal paradigm required subjects to learn the reward value of colors assigned to two stimuli. The color value remained constant for blocks of at least 30 trials before un-cued reversals (**Fig. 1a**). During each trial the onset of color served as the *Attention Color Cue* for shifting covert attention to the stimulus with the rewarded color, while an *Action Motion Cue* was shown before or after the Attention Cue to associate a saccade direction (up- or downwards) with the selection of each stimulus (**Fig. 1b**). After both cues were shown, the animals had to detect a Go-signal (a transient dimming) in the stimulus with the rewarded color to make a saccade for reward. Similar events of the stimulus with the nonrewarded color needed to be ignored and thus served as distracting No-Go events. Monkeys learned the newly rewarded target object in each reversal block within 11 (SE: 2) trials showing learning curves that separated a learning period from a learned period with asymptotic probability for making rewarded choices (**Fig. 1c**).

**Fig. 1.**
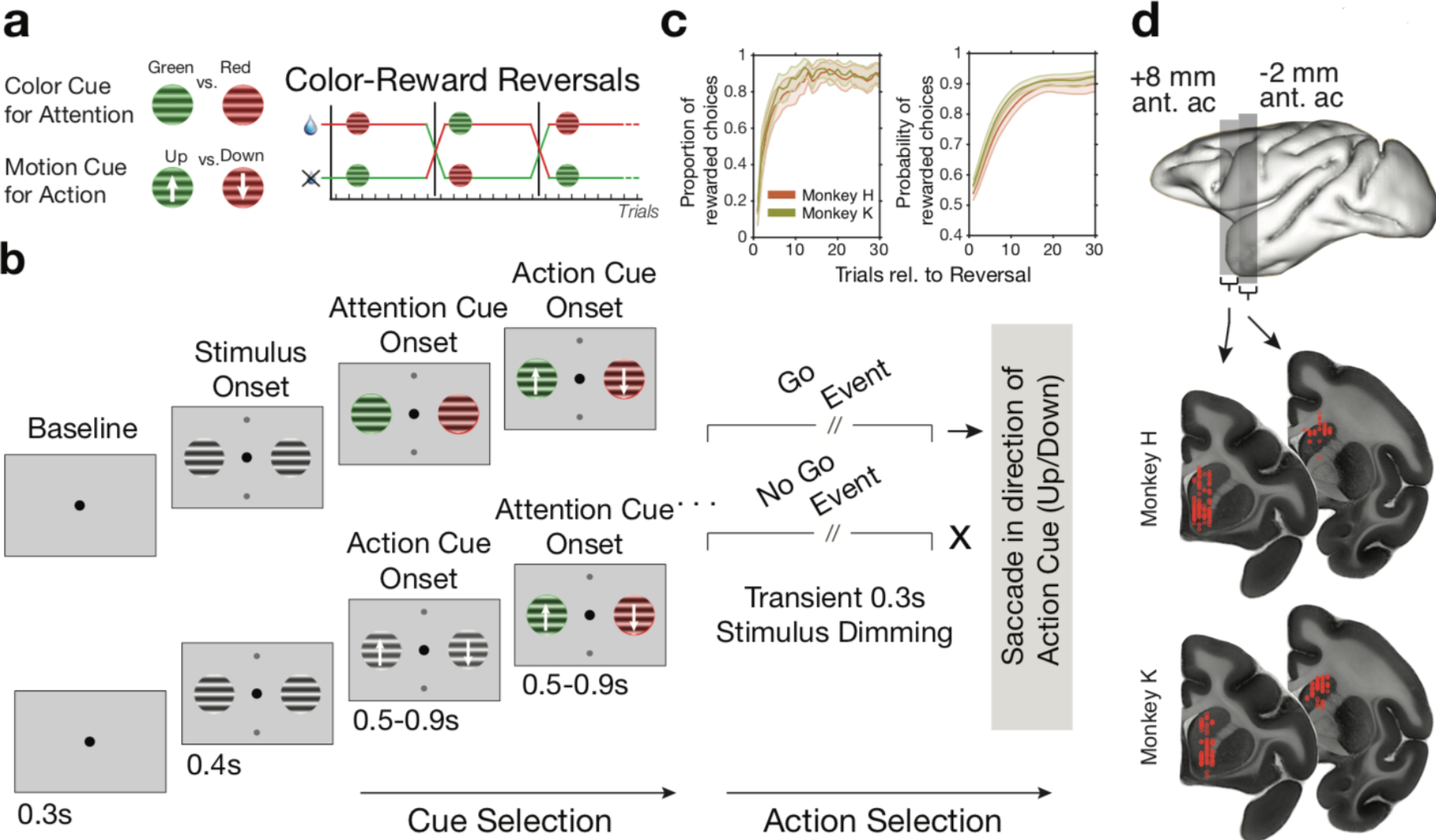
Task paradigm **(a)** The animals performed a color-based attentional learning task requiring learning to attend to the stimulus with the rewarded color and make a saccade in the direction of the motion of that stimulus. The *Color Cue* for attention and the *Motion Cue* for (saccadic) action varied independently from one another. Learning which color is rewarded proceeded in blocks of 30 trials with uncued color-reward reversals. (**b**) During a trial animals fixated the center and were shown the *Attention Color Cue* or the *Action Motion Cue* first. After 0.5-0.9s the remaining cue was shown. Each stimulus could then transiently dim. The dimming was either in the target stimulus first, the distractor stimulus first, or in both stimuli simultaneously. The animal could make a saccade in the 0.05-0.55 s after dimming to receive feedback (reward or no reward). Since only one stimulus was rewarded per trial a dimming event was either a *Go-Cue* to make a saccadic eye movement when it occurred in the rewarded stimulus, or it was a *No-Go Cue* for withholding a movement when it occurred in the non-rewarded stimulus. **(c)** Both monkeys learned the color-reversal task reaching ~80% plateau performance (*left*). We estimated the learning status of the animals with an ideal observer statistic for the probability to observe rewarded choice (*right*). (**d**) Recording locations in the anterior striatum for monkey H and K.

While animals were flexibly performing reversal learning we recorded from 350 neurons in the anterior striatum of two monkeys (164/186 in Monkey’s K/H, **Fig. 1d**). Neurons fell into three separate action potential classes showing broad (B), medium (M), and narrow (N) spikes (**Fig. 2a**) (three-Gaussian model, p<0.01, **Fig. S1a-c**). This tripartite split is similar to prior studies in rodent striatum showing that spiny projection neurons (SPNs) and fast spiking striatal interneurons (FSIs) have broad and narrow spikes, respectively [19–21]. FSIs and SPNs are also distinguished by how bursty and regular their firing is [22, 23], which we quantified using their spiketrains’ *local variability* (LV, [24]) and *global variability* (coefficient of variation, CV) (**Fig. S1d,e**). We combined these firing parameters with the action potential parameters (rise and decay times) for improving the separation of FSIs using cluster analysis. We found that firing patterns and action potentials explained 93.9% of the dataset variance and distinguished two narrow spiking cell classes (N1, N2), four broad spiking cell classes (B1-4), and the class with medium action potential width (M1) (**Fig. 2b,c**, **Fig. S1f,g**). Inspection of the spike rasters (**Fig. 2c**) and spike parameters suggests apparent mappings of these cell classes to classes defined using molecular tools and morphology variables (see **Supplementary Text**). N2 and M1 fulfill criteria of FSIs by showing high firing rates and repetitive interspike intervals (LV’s significantly smaller than 1, random permutation test, p<0.01, **Fig. S1h**). Low firing rates and narrow spikes associate the N1 class with low threshold spiking cells (LTS). The broad spiking classes B1-B4 are likely dominated by SPNs, while among these B classes B1 and B4 will likely include subsets of cholinergic interneurons and neurogliaform cells (see **Supplementary Text**).

**Fig. 2.**
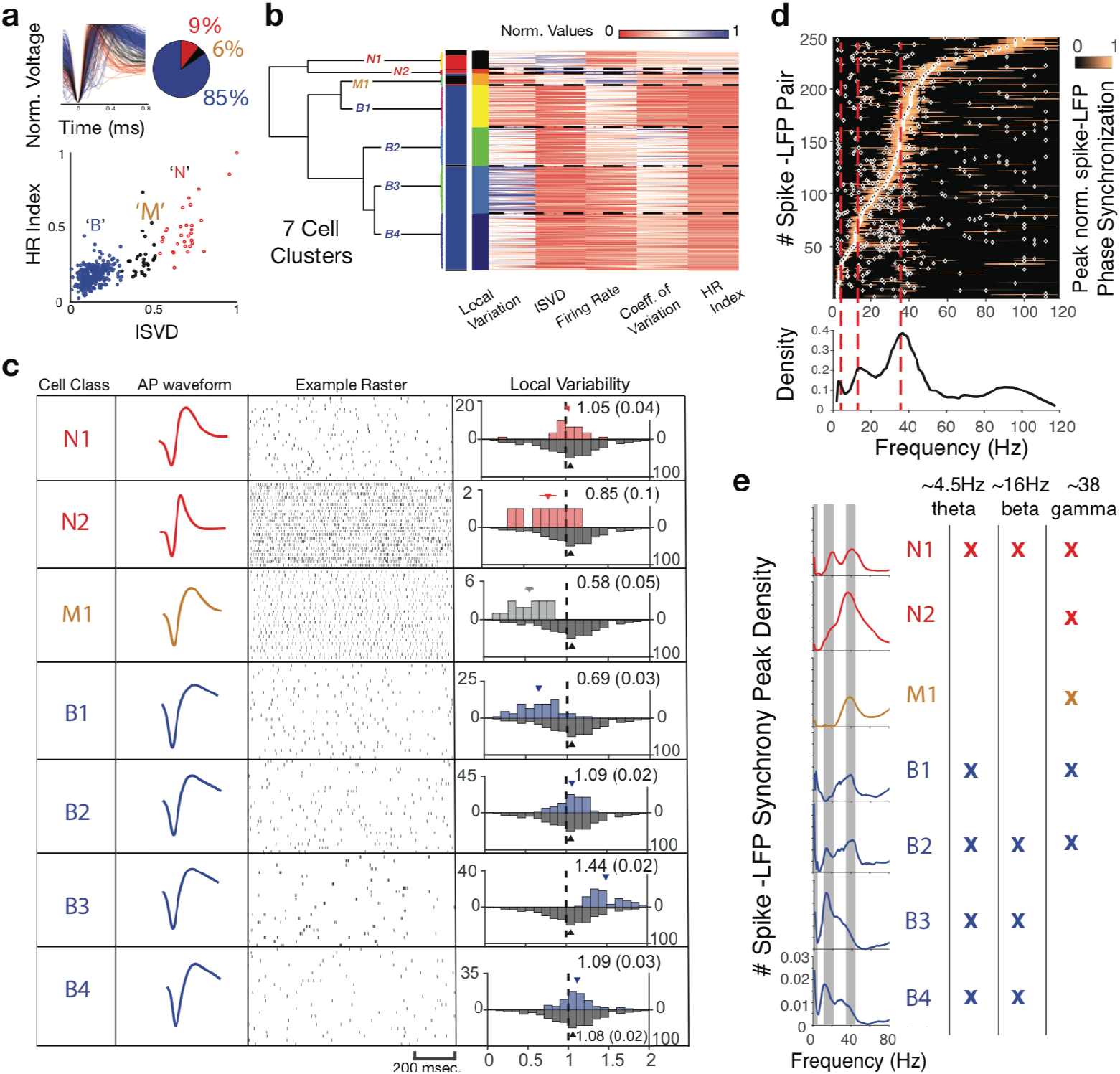
Distinguishing striatal interneurons and spiny projection neurons. (**a**) Normalized, average action potentials (APs) of striatal cells, color coded into 85% broad (B), 6% intermediate narrow (M), and 9% narrow spiking neurons. These three AP classes are distinguished by their Initial Slope to Valley Decay (ISVD), and their Hyperpolarization Rate (HR) (**Fig. S1a-c**). **(b)** Dendogram showing that seven cell classes are distinguishable with unsupervised k-means clustering taking into account the cell’s ISVD, HR index, firing rate, local variability (LV), and coefficient of variation (CV). The clustering identifies four B classes, two N classes and one M class (**Fig. S1d-g**). **(c)** Example rasters, average AP waveforms, and distribution of LV’s for each cell class. Three neuron classes showed highly regular inter spike interval distributions (N2, M, and B1), B3 showed high burst firing propensity, and the other three classes (N1, B2, and B4) showed low firing rates and variable CV’s (**Fig. S1h**). **(d)** Peak normalized spike-to-local field potential (LFP) synchronization for all cells (*y-axis*) across frequencies (x-axis). Synchrony peaks across cells occurred most likely around 4.5 Hz (theta), 16 Hz (low beta), and 38 Hz (gamma) frequencies. White dots indicate significant (p<0.05) synchrony peaks. (**e**) Density of significant synchrony peaks shows that theta and beta synchrony was more likely in broad spiking cell classes B1-4, while gamma band synchrony was more likely in non broad spiking classes N1, N2, M1 (see **Fig. S3**).

In addition to class-specific firing patterns, cell classes also synchronized differently to local striatal field potentials (*see* **Methods**, **Fig S2** and **S3**). We found that cells synchronized significantly to either theta (4.5±3 Hz), beta (16±8 Hz), or gamma (38±10 Hz) frequencies (**Fig. 2d**). The FSI classes (N2, M1) showed prominent gamma band synchronization, but no beta frequency synchrony peaks, while the SPN associated classes B2-4 synchronized prominently at the beta frequency band (**Fig. 2e**). Using support vector machine classification, we could predict for most cells their class label from the cells’ strength of spike-LFP synchronization (**Fig. S3d**). This finding showed that putative SPN classes showed class specific beta band synchrony, while putative FSI classes M1 and N2 showed gamma synchrony consistent with evidence from the rodent striatum [19, 25, 26].

Distinguishing FSIs (M1, N2) from SPN dominated cell classes allowed us testing how these neuron classes responded to the attention cue onset. We found in multiple examples that striatal neurons responded strongly to the onset of the attention color cue, regardless of whether it appeared before or after the action (motion) cue during a trial (**Fig. 3a**, for more examples **Fig. S4**). Across the population, a majority of striatal neurons showed maximal firing rate modulations in the 0.4 sec after attention cue onset (**Fig. 3b**), while response modulations to the action motion cue were rarer and less pronounced (**Fig. 3c**). This attention cue specific responding varied between cell classes. One FSI class (N2) showed a significant, fast and transient on-response to the attention cue (**Fig. 3d**, p<0.05 increase from 0-0.1 sec after cue onset, random permutation test), that was not evident for the action cue (**Fig. 3e**). When N2 cells ceased firing cells of the B2, B4, and N1 class began to show increased firing to the attention cue (Fig. 4d, p<0.05, see **Fig. 3d**) but not to the action cue (**Fig. 3e**). In contrast to these firing increases, fast spiking M1 interneurons reduced their firing to the attention cue (**Fig. 3d**, p<0.05 decrease from 0.2 sec after cue onset onwards, random permutation test). Direct comparison of attention versus action cue onset responses confirmed that N1, N2, M1, B2, and B4 showed stronger modulations to the attention cue than the action cue onset (**Fig. 3e**). These cell-specific attention cue responses were seen irrespective of whether the rewarded stimulus was ipsi- or contralateral to the recorded hemisphere (**Fig. S5**).

**Fig. 3.**
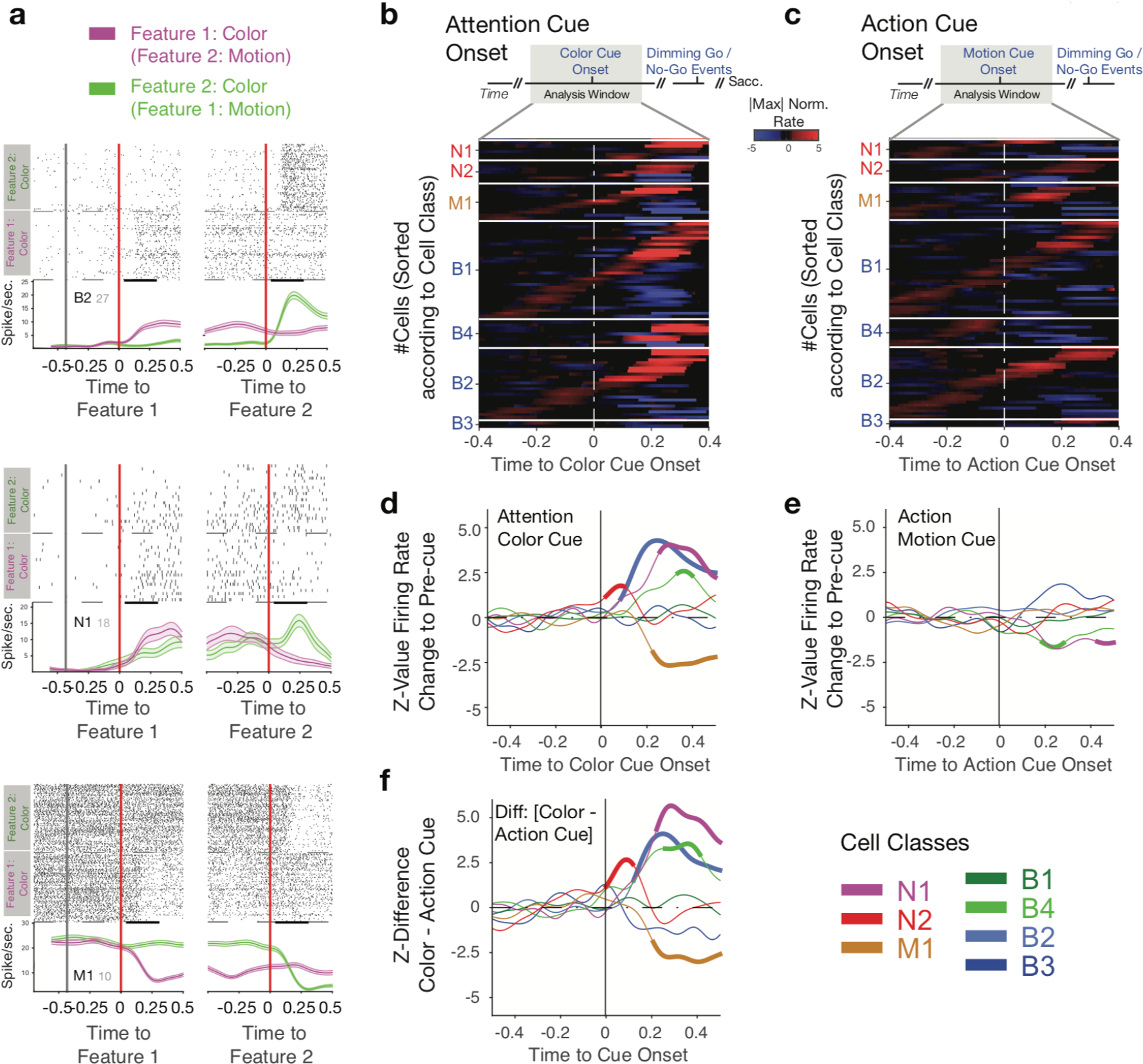
Cell class specific firing rate changes to the Attention Color Cue Onset. (**a**) Three example cells with spike raster around onset of the first cue (*left*) and second cue (*right*). The cells changed firing selectively when the Attention Color Cue was onset either as first cue in the trial (purple), or as second cue (green). They showed no, or less, modulation to the Action Cue onset. Examples are from classes B2, N1, and M1. (**b**) Striatal cells (y-axis) responded to the attention cue onset (0s on x-axis) as shown by normalizing firing to the maximal or minimal firing around the time of Attention Cue onset. (**c**) There were less neurons responding to the Action Cue onset compared to the *b*. (**d**) Z-normalized, average firing around the Attention Color Cue onset show that classes N1,N2, B2, and B4 showed periods with significant firing increase to the Attention Cue, while class M1 shows on average suppressed firing. (**e**) There was markedly lower average modulation to the Action cue. Only N1 and B4 had brief periods of significantly suppressed Action Cue firing. (**f**) The difference of firing to the Attention Color Cue versus Action Cue across cell classes. Thick lines indicate significance (p<0.05, randomization test).

**Fig. 4.**
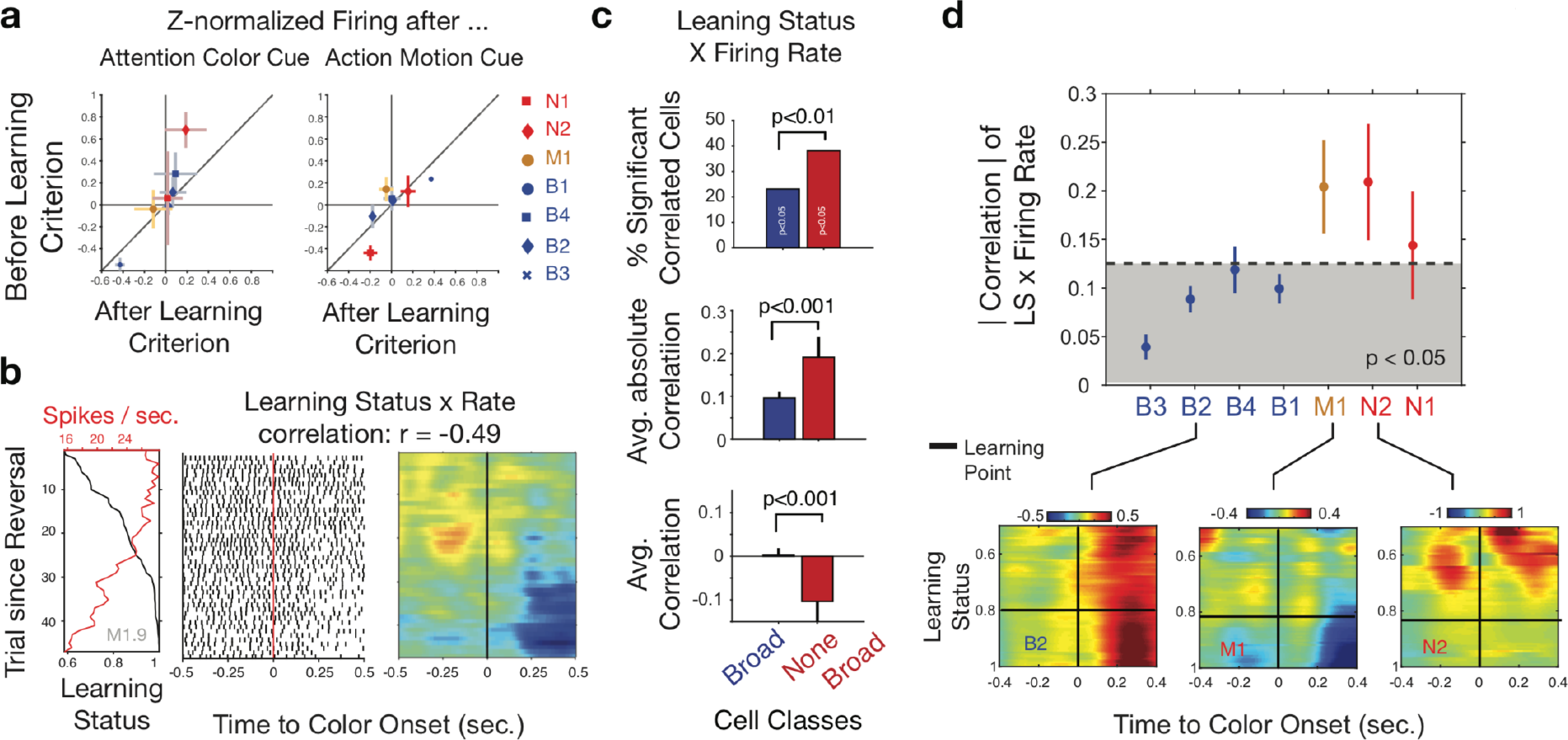
Attention Cue Onset responses change with behavioral learning. **(a)** Average Z-normalized firing rates in a 0.3s time window after Attention Cue for trials before (*y-axis*) and after (*x-axis*) animals reached learning criterion. Errors denote SE. **(b)** Left panel: A single reversal block from an example cell shows gradually reduced firing (red axis) with increased probability of rewarded choices (learning status, black axis). Spike raster (middle panel) and firing rate heat map (right panel) illustrate that this cell shows negative rate x learning correlations (r = −0.49, p<0.01). **(c)** Percent of cells with significant rate x learning correlations (*upper*), avg. absolute correlation strength (*middle*) and avg. correlation (*bottom*). None-broad cell classes show significsantly *more* and *stronger* correlations, that are on average significantly *negative*. (**d**) Absolute correlation of learning status (probability of rewarded choices) and Z-normalized firing rate across cell classes to the Attention Color Cue. Grey shaded area denotes correlation range expected irrespective of cell class label (randomization test). Color panels (*bottom*) show average learning x rate correlations for class B2, M1, and N2. See **Fig. S6a-c** for more details and for correlations after partializing out the expected value of the relevant color and the choice probability.

We next tested how the attention cue responses varied with the reversal learning of color values. We estimated the learning progress as the increased probability of subjects to make rewarded choices, which we call Learning Status (*LS*) (**Fig. 1c**) and used as learning criterion the trial when the lower confidence bound of the probability surpassed chance level [27, 28]. First, we found that the average firing prior to reaching the learning criterion tended to be larger for most cell classes for the attention cue epoch but not the action cue epoch (**Fig. 4a**). To understand this learning relation further we correlated for each cell the post-cue firing rates with behavioral learning (with the learning status, *LS*). Neurons with significant LSxRate correlations show rate modulation that vary systematically with improved color-reward learning such as the example cell in **Fig. 4b** with gradually reduced firing as the animal learns the new color rule. Across the striatum we found that non broad spiking neurons (M1, N1, N2) compared to broad spiking neurons (B1-4) were more likely to show significant LSxRate correlations (38% vs 22%, p<0.01, χ^2^ test), showed stronger absolute average correlations with learning (r=0.19 vs r=0.09, p<0.001, bootstrap randomization test), and showed on average significantly negative correlations (r= −0.10 vs r=0.00, p<0.001, bootstrap randomization test) (**Fig. 4c**). None-broad spiking cells were more likely to fire stronger early during learning or were suppressed after learning had occurred. Importantly, among the none-broad spiking classes, it was particularly the fast spiking neuron classes M1 and N2 that showed significantly stronger correlation compared to other classes (**Fig. 4d**). Both these neuron classes showed on average negative correlations (**Fig S6a**).

The correlation results so far were between firing rates and an ideal observer estimate of reward reversal learning that can be interpreted as the degree of confidence of the subject that a specific color leads to reward [27]. Concomitant to increased confidence about which color is relevant, learning is also accompanied by increased expected stimulus values and increased choice probability (**Fig. S6b**), raising the question which learning variables best correlates with neural firing. We estimated choice probability (CP) and value expectations (EV) with a reinforcement learning model (see **Methods**) and found that both, CP and EV also correlated with firing rates, but with partial correlations only the ideal observer estimated confidence estimate remained significantly negatively correlated with none-broad spiking cell firing (**Fig. S6d**). These findings indicate that striatal FSI firing to the attention cue is closely linked to an internal state of confidence about the behaviorally relevant color feature.

The firing changes to the attention cue could reflect the gating of the rewarded target stimulus, or the suppression of the non-rewarded distracting stimulus. To disentangle these scenarios, we compared the responses of neurons to the transient dimming events that could occur only in the rewarded target stimulus (Go event), or only in the non-rewarded distractor stimulus (No Go event). We found multiple example neurons that selectively responded to the Go event by increasing or decreasing their firing (**Fig. S7a**) and confirmed that on average five of seven cell classes (N2, M1, B1, B2 and B4) showed significant responses to the Go event indicating the gating of the target (**Fig. S7b**). However, we likewise found that three cell classes (M1, B3, and B4) significantly increased activity to the No Go distractor event indicating that subsets of cells increased their firing to a salient distractor event that needed to be suppressed (**Fig. S7c**). These findings highlight that both, attentional gating of relevant cues and filtering of salient distractors are represented in cell class specific firing increases. Statistically, the gating responses were stronger than the filtering responses (**Fig. S7d**). These gating and filtering responses were changing with behavioral learning progress for classes M1 and B4 (Pearson correlations, p<0.01, multiple comparison corrected, **Fig 7e**), and for class M1 the Go and No-responses correlated on a trial-by-trial basis with the onset responses to the Attention Color Cue (Dunn’s test, p <0.05, **Fig S8**). These findings highlight that the FSI’s of the M1 class gate events not only when they direct covert attention (to the color onset), but also when they promote overt (saccadic) choice behavior (to the Go event).

The observed cell specific correlations show that two separate classes of putative FSI’s change their firing to the sensory color information that cues selective attention. One FSI class (N2) showed strongest activity during learning indicative of stronger inhibition when new reward predictions are formed. The other FSI class (M1) showed average reduced inhibition after reward learning occurred indicative of disinhibition of spiny projection neuron responses. These cell type specific cue responses were closely associated with the gating of target information (Go) and linked to the subjects confidence about the reward value of stimuli reminiscent of an *attentional policy* that determines how much attentional resources should be allocated to different visual features (here: colors) [29, 30]. When visual features are presented, such a policy could shift covert attention to the most relevant visual stimulus. We found that such striatal mediated attentional shift involves both, the enhanced throughput of target changes as well as the active suppression of distractor changes akin to Go and No Go responses in the motor domains facilitating a target action and inhibiting conflicting actions respectively [15, 31].

In summary, our results suggest a critical role of the striatum and its inhibitory circuits for the learning of feature based attention biases that underlie object choice [7, 31].

## Acknowledgments

The authors would like to thank Stephanie Westendorff for help with the design of the study.

## Funding

This work was supported by a grant the Canadian Institutes of Health Research MOP 102482 (T.W.). The funders had no role in study design, data collection and analysis, the decision to publish, or the preparation of this manuscript.

## Author contributions

T.W. conceived the experiment. M.O. and S.A.H, performed the experiments. K. B.B., and T.W. analyzed the data. T.W. and K.B.B. wrote the original draft. All authors edited the paper;

## Competing interest

The authors declare no competing interests.

